# Thermal performance in fishes varies systematically across latitude, habitat, and biological organization

**DOI:** 10.64898/2026.03.05.709660

**Authors:** Hannah Mosca, Nikki Moore, Laura Gervais, Jennifer Sunday

## Abstract

Thermal performance curves are widely used to understand how ectotherms respond to anthropogenic climate change, yet comparable data remain fragmented for fishes despite their predicted sensitivity to warming. Here we introduce FishTherm, a compilation of 457 thermal responses from 107 wild fish species, spanning diverse performance metrics and temperatures. With this dataset, we estimated thermal optima, performance breadth, tolerance breadth, and activation energies, and tested how these thermal traits vary geographically, across latitude and habitat, and by type and functional context of performance. Thermal optima increase toward lower latitudes and warmer waters, consistent with established biogeographic patterns. We also found that responses measured at lower levels of biological organization exhibit higher thermal optima than population-level responses, supporting complexity constraints on thermal performance curves. Activation energies broadly matched predictions from the Metabolic Theory of Ecology, and we further found systematic variation of lower values in negatively-motivated responses, and associated with narrower thermal performance breadths, in line with theory. Our results provide supporting evidence for several unifying principles that advance our understanding of constraints and variation in thermal performance. FishTherm is a new resource for comparative thermal ecology and for improving our understanding of fish vulnerability under climate warming.

## Introduction

Increases in global average temperature combined with changes in the frequency and duration of extreme heat events are exposing organisms to new climatic conditions [1]. In response, species ranges are moving rapidly across latitude, elevation, and depth [2]. As ongoing shifts in temperature regimes restructure ecosystems and affect human-relevant services [3], understanding species’ physiological responses to temperature may be a useful approach to predicting future change under anthropogenic warming.

Among ectothermic organisms, whose body temperature closely tracks ambient conditions, environmental temperature governs the biochemical and physiological processes that drive life [4–6]. A common method to evaluate the physiological effects of temperature examines how body temperature affects performance, an organism’s ability to function within its environment, assessed by measuring ecologically-relevant variables, like swimming speed, metabolic rates, growth rates, and even overall fitness [7]. When these processes are measured over a range of body temperatures, they often follow a unimodal curve known as a thermal performance curve (TPC), where performance increases to an optimal temperature (*T*_*opt*_ ) and subsequently declines sharply due to physiological stress or failure [8–10]. We can extract key parameters from TPCs describing thermal sensitivity of organisms [11,12], including thermal optima (*T*_*opt*_ ), upper and lower performance limits, ‘pejus’ temperature (where performance begins to decline [13]), various metrics of performance breadth (e.g. thermal tolerance breadth, breadth of 80% of maximum), and the rate of performance increase towards the optimum. This rising limb reflects the activation energy of a Boltzmann–Arrhenius function, consistent with the temperature dependence of underlying metabolic processes [5,14–16].

Thermal performance curves often align – either through evolutionary adaptation or habitat selection – with thermal environments experienced by individuals or their populations. As such, the temperature at maximum performance (i.e. thermal optimum) and maximum temperature of non-zero performance (i.e. thermal maximum) is often greater in species that experience warmer temperatures [17–19]. Further, in more variable environments, not only does breadth of performance tend to be wider to accommodate fluctuations in temperature, but thermal optima tend to be cooler than mean environmental temperatures compared to species in less variable environments [20]. Indeed, terrestrial insects in the tropics experience temperatures closer to their thermal optima than higher-latitude species [18]. In more variable thermal environments, the fitness optimum is expected to be lower than that measured under constant conditions, because temperature fluctuations around a ‘concave’ peak disproportionately reduce long-term average performance near the upper thermal limit. We therefore expect selection to lead to a higher thermal optima in more variable conditions [21]. It is also thought that species with greater capacity for behavioural thermal regulation might demonstrate TPC shapes that are less reflective of environmental temperatures, as behaviours can reduce the necessity of physiological adaptation [22].

Considering the rising portion of a performance curve, the Metabolic Theory of Ecology predicts a universal temperature dependence of biological rates [5,14,15], yet variation in thermal sensitivity exists and could reveal important mechanisms. In a broad compilation of activation energies across diverse biological responses and taxa, Dell et al. found evidence that the functional setting of a response might alter its temperature dependence [16], advancing a hypothesis consistent with the ‘life–dinner principle’ – the expectation that selection is stronger for survival or escape from death than for marginal increases in energy acquisition [23,24]. Under this hypothesis, Dell et al. expected negatively-motivated processes such as prey escape and defense would show reduced temperature sensitivity because these mechanisms would be under strong selection even at sub-optimal temperatures. Indeed, they found that among many organisms, with the majority being insects, performance increased more gradually per unit temperature (lower activation energy) for responses to negative stimuli (e.g. being chased) than for positively-motivated (seeking food), or autonomic processes (resting metabolic rate)[16].

Structural constraints to the shape of TPCs could also drive variation in thermal dependency. Thermal optima and activation energies are both hypothesized to change systematically with the complexity of the biological response, due to combinatorial effects of multiple rate-limiting processes at higher levels of organization [25,7,26]. Indeed, among biological processes of Drosophila, higher-order processes had decreased thermal optima and steeper activation energies, in support of the complexity-shifted-TPC hypothesis [27]. Taking this a step further, if activation energies are variable among species and response types, another constraint could be a trade-off between the width of the curve and the height (i.e. a thermal specialist/generalist trade-off [28]). Under this hypothesis, the slope of thermal sensitivity would be shallower when thermal performance curves are broader [29]. This has been detected in a global analysis of fish thermal tolerance limits, in which broadly-tolerant eurythermal organisms were found to have lower activation energies [30].

Compiling thermal performance curves and testing hypotheses about variation in their parameters can improve our understanding of the evolutionary drivers and constraints that shape thermal performance and provide new integrated information for estimating vulnerabilities to climate change [7,31]. Studies of thermal tolerance limits (*CT*_*min*_/*CT*_*max*_) suggest that thermal tolerance breadths of aquatic species match their experienced temperatures [32] and track climate changed more closely [33] compared to terrestrial species, and that they vary by life history stage [30], underscoring the heightened vulnerability of aquatic organisms to warming [34]. Although aggregated datasets of full thermal performance curves exist for other organisms, e.g. phytoplankton [17], reptiles [35,36], and insects [18], a compiled dataset for fish is lacking. A compilation of TPCs across many taxa was the subject of an impressive comparative analysis of TPC curve parameters [16], but these appear to represent multiple responses from only 11 fish species (based on datasets available in Kontoupalous et al.[37]). Dalke et al. compiled thermal responsiveness from 130 fish species, but these data were limited only to the rising portion of the curve and included only development and metabolic rates [30]. Because fish span both marine and freshwater habitats, a global compilation of a multitude of response types can allow us to test for relationships between environmental variability and thermal sensitivity [38]. Given their central role in food security and economic stability, understanding the responsiveness of fish species to climate warming is a critical priority [39,40].

To fill this gap, we conducted a systematic quantitative review and data extraction of thermal performance curves for fish, and used the resultant dataset to test hypotheses about variation in thermal performance. We included thermal responses of freshwater and marine fishes, codified the methods, geographic origin, acclimation conditions, and other metrics of each response assay, and fit appropriate models to characterize curve shapes and estimate curve parameters to each assay. We used this dataset to test how fish thermal optima vary with latitude and environmental temperatures. Under the hypothesis that species are adapted to their experienced temperatures, we expected *T*_*opt*_ to decrease with latitude and increase with environmental temperature, with variation falling around a 1:1 relationship. According to the hypothesis that thermal optima are selected to be warmer than mean conditions in more variable environments (Suboptimal-is-optimal) [21], we also expected thermal optima to be greater compared to mean environmental temperatures (greater difference, i.e. thermal safety margin) in fish that experience more temporal-variability in temperature. Thus, we expected a greater thermal safety margin in freshwater vs. marine, and increasing thermal safety margin with increased temporal variability. According to the complexity-shifting-TPC hypothesis, we expected lower thermal optima in biological responses at higher orders of complexity (population>organismal>internal [16]). Considering the rising portion of TPCs, we also assessed the central tendency of activation energies against the expectation of 0.65eV as the fundamental metabolic activation energy, and the hypotheses the activation energies would be lower in more negatively-motivated (“escape”-type) responses, at lower levels of biological organization (15), and in wider TPCs [30]. Our results advance our understanding of constraints and variation in TPCs, and we expect the FishTherm database to be useful in many future analyses.

## Methods

### Literature search and data extraction

We performed a literature search to identify lab-based experimental studies examining the effects of temperature on the performance of wild fish. To decide on a set of search terms and article eligibility criteria, we first conducted a pilot search and article screening. The week of June 9, 2024, we searched the literature using 17 different sets of test search terms designed to retrieve temperature responses across a broad array of both physiological and ecological performance metrics. We tested each term set across four major databases - Web of Science, Scopus, ProQuest, and Lens - without filtering articles based on publication date. For each test search, we scanned the first page of retrieved articles and selected the pilot set of search terms as the test set that maximized the number of articles retrieved without jeopardizing relevance (see Supplementary Methods for terms). To begin our pilot screening, we uploaded all articles retrieved by the pilot search terms from all 4 platforms (48,992 unique records) to Rayyan, an online platform for systematic reviews [41].

We then conducted a pilot screening using a random subset of 10% of the records (approximately 4,320 articles) to refine article eligibility criteria and standardize the screening process. Two reviewers independently reviewed each abstract, categorized article inclusion as either ‘yes’, ‘no’, or ‘maybe’, and tagged articles with corresponding inclusion or exclusion criteria. We then collaboratively resolved conflicts between reviewers, which led to clearer article eligibility criteria. The pilot screening returned 239 relevant articles (5% of total articles screened), indicating that the search terms were too broad. We then refined the search terms to improve specificity while ensuring searches retrieved the relevant articles identified during the pilot phase. Using the final set of search terms selected, two relevant studies from the pilot search were not retrieved. To ensure their inclusion, we added these studies to a list that would later be manually added to the dataset (following methods of Poittier et al. [42]).

We conducted our search using the final set of search terms on September 16, 2024, using the Web of Science Core Collection, selected for its rigorous indexing of high-impact papers [43]. We then screened articles as outlined in the PRISMA diagram (Figure 1). After removing duplicated articles (n = 5), the Web of Science search retrieved 8,854 unique articles. Titles, abstracts, and keywords of these articles were screened in Rayyan following the exclusion criteria developed during the pilot screening, yielding 677 articles for full-text review. Based on the full-text, studies were then retained if they 1) tested fish species under controlled laboratory conditions, and 2) measured performance traits across at least four distinct temperature treatments spanning a minimum temperature range of 5°C. These thresholds ensured sufficient resolution for generating reliable thermal performance curves [16]. As in the screening process, studies were excluded if they were field-based or observational in nature (e.g. those correlating interannual variation in abundance with temperature), if they tested captive or selectively bred fish, or if they only measured critical thermal limits. We also excluded molecular, in vitro, and theoretical studies, as well as to studies that lacked performance metrics or sufficient temperature treatments (see Figure 1 for details). This process yielded 118 eligible studies that met all inclusion criteria, encompassing data on 107 fish species.

**Figure 1.**
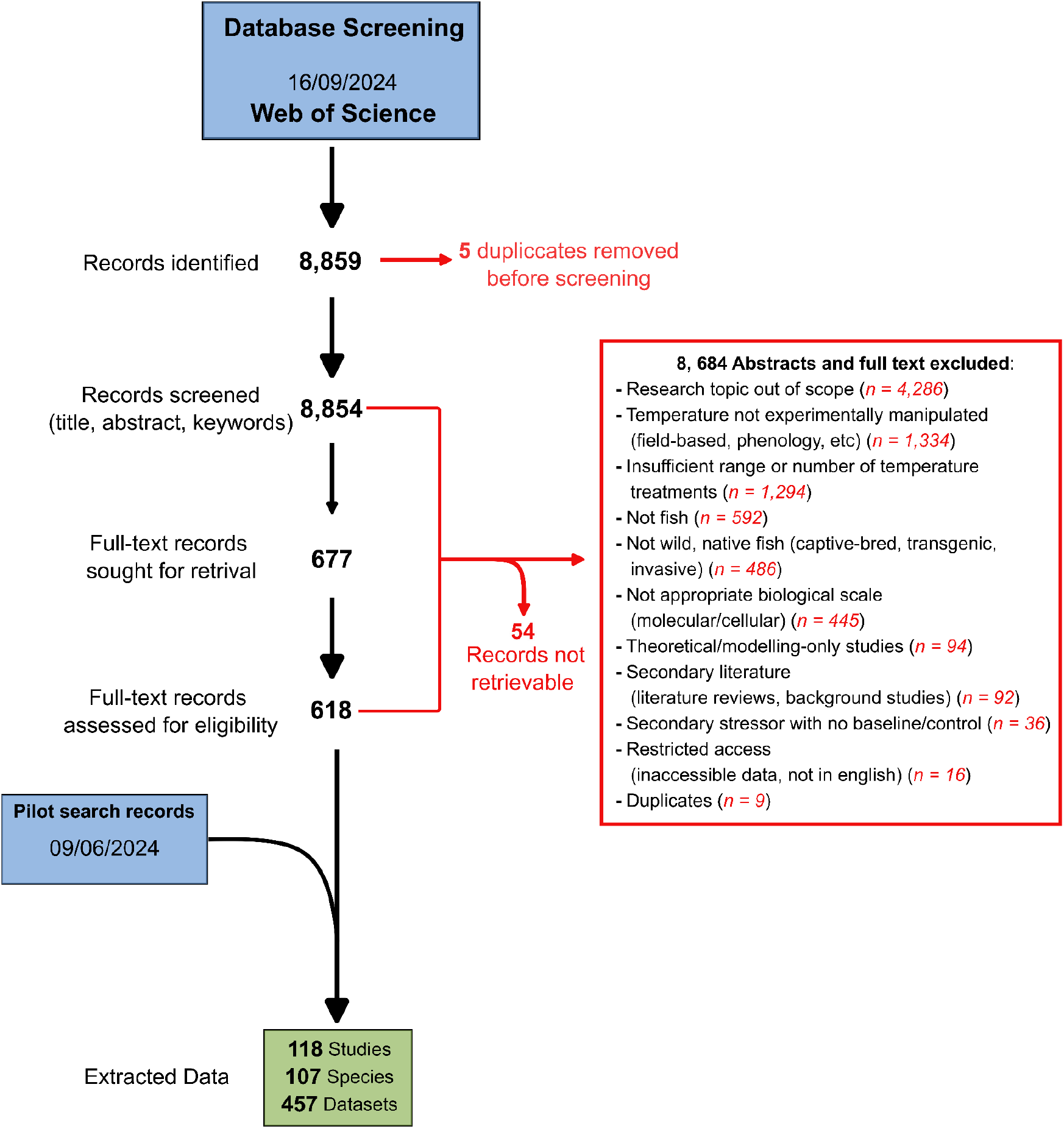
Preferred Reporting Items for Systematic reviews and Meta-Analyses (PRISMA) flow diagram of literature screening and study selection. Flow diagram summarizing the identification, screening, eligibility assessment, and inclusion of studies in the systematic review. Note: Numbers reported in the red box do not reflect the fact that some articles may have met multiple exclusion criteria.

We extracted physiological response data from 118 studies, including mean and individual measurements of performance across temperatures, along with associated sample sizes and measures of dispersion (e.g. standard deviation, standard error). Data presented in the text or tables of articles were extracted directly, while data displayed in figures were digitized using WebPlotDigitizer, a web-based tool designed for extracting data from plots [44]. In addition to measures of performance and temperature, we extracted relevant metadata when reported, including genus, species, age, geographic collection location, habitat type (e.g. freshwater, marine, brackish), body mass, body length, sex, and details regarding the experimental methodology used (e.g. definition of performance metric or holding time). To differentiate between the two general methods used to test thermal performance in a laboratory setting, each dataset was classified as either a ‘batch-acclimated’ or ‘acute’ experiment based on the temperature exposure regime. In batch-acclimated experiments, a group of fish is acclimated to a specific temperature (with acclimation time ranging from days to weeks), and performance is subsequently tested at that same temperature. In contrast, acute exposure experiments involve subjecting fish to a series of test temperatures for short durations (often minutes to hours), without significant prior acclimation to test temperatures. Additionally, we noted sample size when reported, since the number of performance measurements represented by each data point often differed; while most datasets reported mean performance across a group of fish tested, others reported individual-level measurements of performance.

To test whether thermal performance relationships differ among types of biological performance, we categorized each response variable by the type of performance measured, the motivation of the response, and the level of biological organization. Each response was assigned to one of six types: metabolism, somatic growth, locomotion, energy acquisition, reproduction, and survival (Figure 3B). Following the ontology developed by Dell et al., traits were categorized by degree of control, as autonomic (below conscious control), or somatic (expressed through behavior or voluntary movement). Somatic traits were considered to be either negative (escape/defense/maximum oxygen consumption), positive (approach/consumption), or voluntary (discretionary activity not directly linked to defense or consumption, such as activity or routine swimming speed). Also in line with Dell et al., traits were classified at different levels: internal (processes within the organism), individual (whole-organism processes involving interaction with the external environment), interaction (processes involving interactions between species), or population (processes for a group of conspecifics) [16]. Full trait definitions and categorizations are provided in the Github repository.

### Categorizing datasets according to coverage of curves

In initial explorations of the data, we found that the shape and coverage of the full thermal performance curve varied greatly across datasets. While some responses showed a characteristic, left-skewed thermal performance curve, many were not bounded by thermal minima (*T*_*min*_,) and maxima (*T*_*max*_), or were only assayed at the increasing or decreasing portion of the curve (Figure 2B). We therefore grouped datasets using benchmarks to assess the ‘thermal coverage’ of the assay to streamline further analysis.

**Figure 2.**
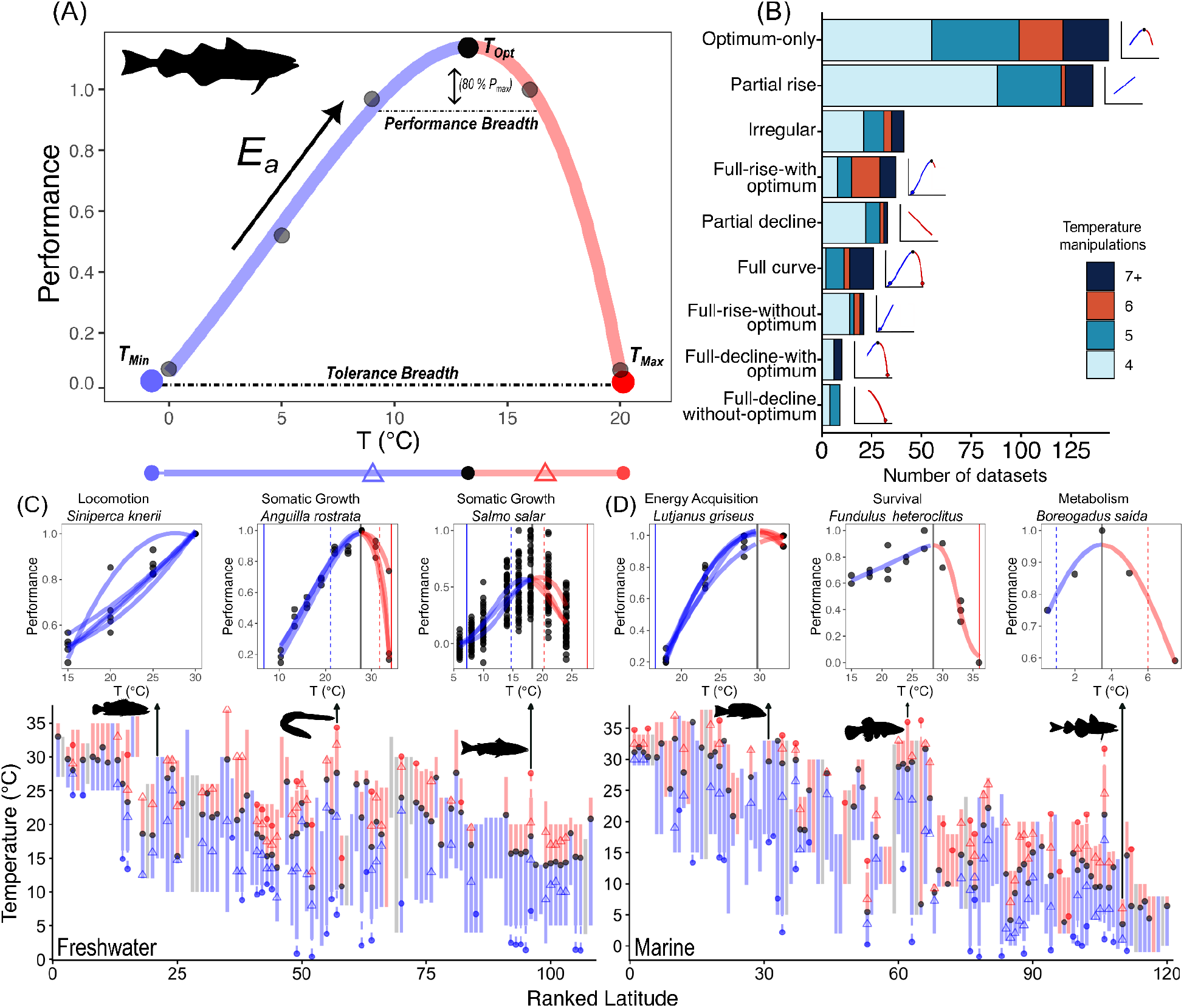
Summary of thermal performance curves (TPCs) for every species, response type, and latitude of collection. (A) Example fitted TPC for *Gadus chalcogrammus* (Walleye Pollock) shown for specific growth rate (% wet weight day^−1^) using the Thomas 2012 model. Small black points represent raw data while large colored points indicate estimated *T*_*min*_, *T*_*max*_, and *T*_*opt*_. The rising slope of the TPC is used to estimate activation energy (E_a_), and horizontal dashed lines show tolerance breadth and performance breadth (temperatures supporting ≥80% of maximum performance). (B) Distribution of datasets categorized by curve-coverage type (see Methods), with bar segments shaded by the number of distinct experimental temperatures used (4 to ≥7). Small schematic curves illustrate each category, with points indicating thermal minima (*T*_*min*_; blue), maxima (*T*_*max*_; red), and optimum (*T*_*opt*_ ; black) when estimable. (C-D) Geographic distribution of thermal traits across freshwater (C) and marine (D) species. Each vertical line represents a ‘top-down’ view of a semi-aggregated TPC (see methods). In panels B-D, the solid lines span the experimental temperature range tested, and dashed lines extend to the extrapolated parameters when outside of the tested range. Line colour indicates increasing (blue) and decreasing (red) portions in the same curve; if only one colour is shown, only a monotonic rise (blue) or decline (red) was observed. Curve parameters, when estimable, are denoted using symbols for *T*_*min*_ (blue point), *T*_*max*_ (red point), *T*_*opt*_ (black point), upper performance breadth (red triangle) and lower performance breadth (blue triangle), as illustrated with curve mapping to horizontal line in panel (A). Insets show six example TPCs expanded to side-view, showing individual curves (red and blue curved lines) and underlying data (black points) that were used for aggregate curve parameters in panels C-D, shown as thin vertical lines to represent *T*_*opt*_ (solid black), *T*_*min*_ (solid blue). *T*_*max*_ (solid red) and upper and lower performance breadth (dashed red and blue).

We first rescaled the response measurements within each thermal performance dataset to enable consistent assessment of response shape across datasets that differ in trait type, and thus response magnitude. For each dataset, we rescaled responses by dividing each response measurement by the absolute value of the maximum observed response within that dataset, preserving true zeros in the original data, resulting in an expression of performance at each temperature relative to the maximum observed response. We then sorted responses into one of nine categories: full curve, optimum-only, full-rise-with-optimum, full-rise-without-optimum, partial-rise, full-decline-with-optimum, full-decline-without-optimum, partial-decline, and irregular responses (Figure 2B). When assigning coverage categories, we defined full curve TPC responses as those exhibiting an increase in performance followed by a decline, with performance approaching zero at both extremes, defined when the rescaled response at the minimum or maximum temperature tested was less than or equal to 0.2 (20% of the curve-specific maximum). We defined optimum-only responses as those exhibiting a single, well-defined performance maximum within the tested temperature range, with rescaled performance increasing monotonically up to a peak and decreasing thereafter. These responses do not approach near-zero performance at one or both thermal extremes and therefore only capture the location of an optimum without fully resolving thermal limits.

Full-rise-with-optimum and full-decline-with-optimum were defined as those that approach near-zero performance at one, but not both, thermal extremes. Compared to full curves, in these cases we had less information on which to estimate thermal limits, and therefore only considered a curve adequate to estimate the cold or warm thermal extreme if it had a rescaled response that was less than or equal to 0.1 (10% of curve-specific maximum). Similarly, full-increase-without-optimum and full-decline-without-optimum were defined as those datasets that captured one of the thermal extremes in this way but without a clear optimum. Partial rise and partial decline responses were defined as those with a monotonic increase or decrease, but without evidence of a performance peak or declining toward near-zero at extremes. Irregular responses were those that fit none of these criteria.

### TPC model fitting and selection

We fit thermal performance curve (TPC) models to each dataset using the R package rTPC [45], a robust framework for fitting nonlinear temperature-performance relationships using a broad set of model formulations derived from the ecological and physiological literature. Within rTPC, candidate models differ in the number of free parameters required to describe the thermal response. We therefore modelled datasets that contained four distinct temperature treatments (our minimum for inclusion, with a minimum spacing of 1°C between temperature treatments), differently than those included five or more distinct temperature treatments. All available three-parameter models were initially fitted to the 220 datasets with four distinct temperature treatments, while all available four-parameter models were initially fitted to the 237 remaining datasets.

We removed four candidate models that were producing consistently poor fits, such as those that forced the data to have multiple local maxima and unsupported shapes in performance curves across temperatures. After screening, we retained two three-parameter models and nine four-parameter models for subsequent model selection. Models were further filtered so that those predicting performance values outside one standard deviation of the observed data were excluded. For bounded responses, where performance approached zero at low or high temperatures, models predicting thermal minima or maxima more than 5 °C beyond the range of tested temperatures were also removed. We selected the top model for each response based on the lowest AIC score, choosing the model that best fit the data among structurally different models with equal complexity, and used it to estimate thermal performance parameters, including *T*_*max*_, *T*_*min*_, *T*_*opt*_, *performance breadth, tolerance breadth*, and *activation energy* when data met eligible criteria.

### Environmental temperature data extraction

To test our hypotheses about how thermal performance varies across environments, we extracted data on the environmental temperatures experienced by fish used in thermal performance experiments. When information on the geographic collection location of tested fish was reported in the original study, we extracted estimates of the temperatures experienced at the collection location from global raster databases of sea and freshwater surface temperature. When only a collection location description was reported by the original study, we estimated point coordinates of collection from that description using Google Maps and then used these coordinates to extract estimates of experienced temperatures.

For marine studies, we extracted sea surface temperature (SST) data from the Environmental Research Division’s Data Access Program (ERDDAP) developed by NOAA’s Southwest Fisheries Science Center. The dataset used was from NOAA Global Coral Bleaching Monitoring SST and SST Anomaly product (Version 3.1, 5 km resolution, monthly, 1982–present), provided by the NOAA/NESDIS/STAR Coral Reef Watch program (Dataset ID:NOAA_DHW_monthly_Lon0360). We extracted mean monthly values from September 1982 through September 2025.

In freshwater studies, estimates of freshwater surface temperature were extracted from FutureStreams, a global dataset of modeled streamflow and water temperature projections at 5 arcminute spatial resolution [46]. We used the E2O reanalysis-forced historical simulation for freshwater conditions for the historical period, and the HadGEM2-ES climate model projection under RCP4.5 to represent contemporary conditions beyond the historical period. We used these to compile a continuous time series of mean monthly temperature values spanning September 1982 through September 2025, matching the time span of the marine SST data, using the E2O simulation through December 2005 and the HadGEM2-ES RCP4.5 simulation thereafter.

From freshwater and marine mean monthly water temperatures, we then calculated three summary metrics to represent the average, hot extreme, and variability in experienced temperatures respectively: (1) monthly mean temperature, calculated as the average mean monthly temperature across all months, (2) upper temperatures, calculated as the upper 97.5 percentile of mean monthly temperature across all months, and (3) thermal variability, calculated as the standard deviation in mean monthly temperature across all months.

### Treatment of pseudoreplicates

Often, a single study reported multiple response types measured from the same individuals or experimental groups. While some responses from the same fish may reasonably be considered independent measures (e.g. growth rate and prey attack rate), often responses were closely related (e.g. consumption rate and conversion efficiency) and therefore could not be considered independent measures. While we report all raw values in the TPC dataset, for analyses and visualization we accounted for non-independence by defining pseudoreplicates as performance responses originating from the same study, measured in the same species, collected from the same location, and belonging to the same response types (i.e. one of six response groups in Figure 3B). This yielded individual estimates of thermal responses from which we could test for differences between performance trait groups (e.g. metabolism vs. swimming speed) while reducing pseudoreplication within groups [16] (See insets in Figure 2C and D).

**Figure 3.**
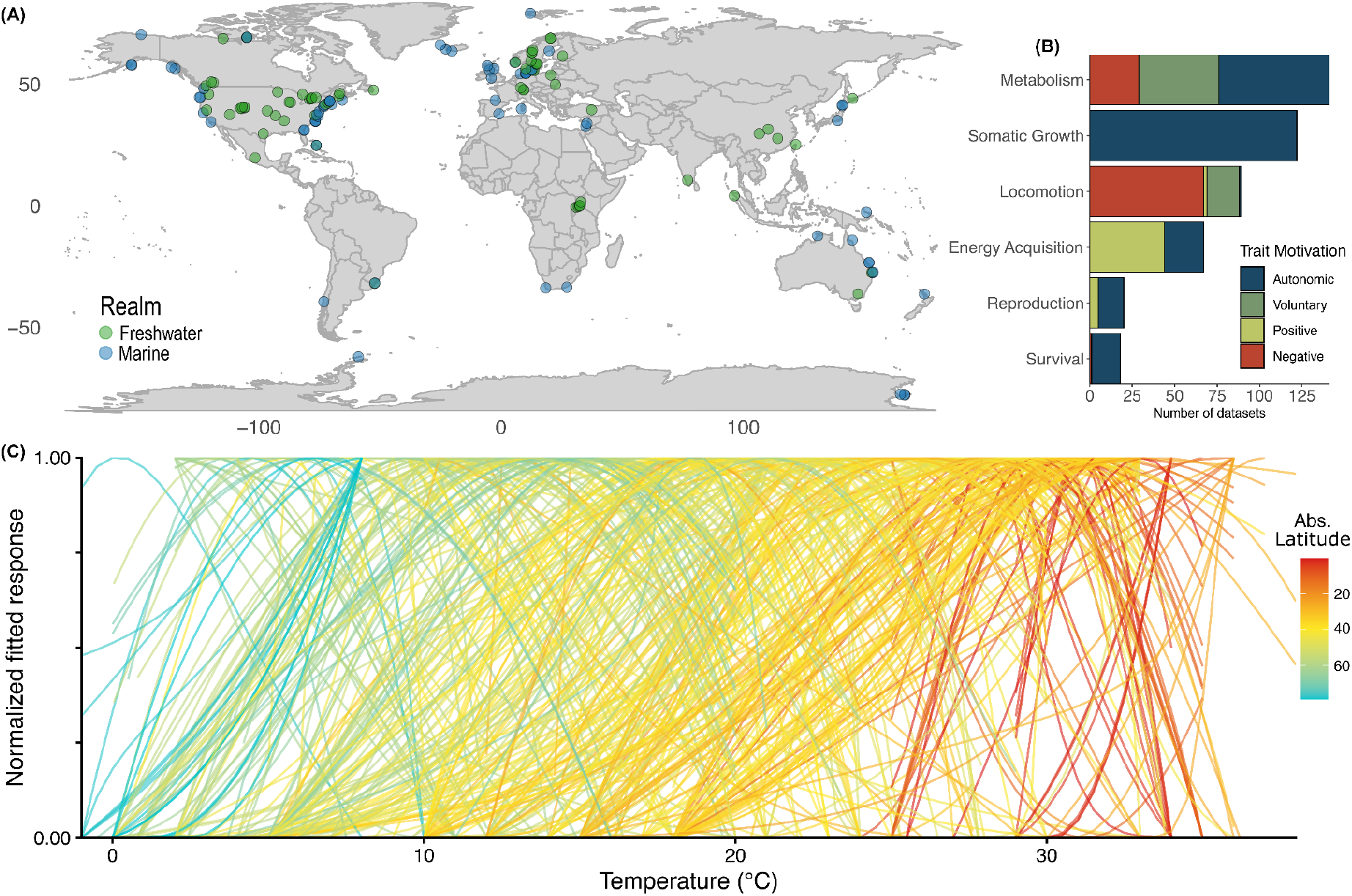
Geographic distribution and trait composition of thermal performance data. (A) Global map of fish collection locations extracted from eligible studies, colored by realm (marine vs. freshwater). (B). Distribution of physiological performance traits measured, with bars subdivided by trait motivation (autonomic, voluntary, positive, and negative). (C) Predicted thermal performance curves across all dataset types, including bounded and unbounded responses. Model-fitted responses were normalized within each curve using min–max scaling (0-1), allowing comparison of thermal response shape and positioning independent of response magnitude. Line color represents absolute latitude.

For analyses that further separated responses by motivation and level of biological organization, pseudoreplicates were further grouped by motivation and organization level. This resulted in a finer classification, such that responses that were previously aggregated within the same response type group (e.g. resting and maximum ventilation rates) were treated as separate response categories, allowing us to test whether functional response motivation and level of biological organization influence activation energy and thermal optima.

For each pseudoreplicate group, we obtained a single value of each curve parameter (*T*_*max*_, *T*_*min*_, *T*_*opt*_, *performance breadth, tolerance breadth*, and *activation energy*) by averaging parameter estimates across eligible responses within that group. However, these resulting semi-aggregated curves still carried non-independence (e.g. multiple semi-aggregated curves could result from the same study or even the same individuals when different performance trait groups were considered), which was further accounted for in our modelling by holding ‘study’ as a random effect.

### Testing hypotheses

To test our hypotheses regarding how thermal optima (*T*_*opt*_ ) vary with latitude and environmental temperature, we fit a set of linear mixed-effects models to *T*_*opt*_ . We ran three separate models that included either (1) absolute latitude of collection location, (2) monthly mean water temperature at collection location, or (3) upper water temperature at collection location as the primary fixed-effect predictor. To evaluate whether thermal performance breadth and thermal safety margins vary with environmental temperature variability, we fit two additional linear mixed-effects models using the standard deviation of monthly mean water temperature as a fixed effect. In all models, we included an interaction between the environmental predictor and realm (freshwater vs. marine) to test whether these relationships differ between habitat types (for the purpose of analysis, species collected from brackish habitats were grouped into either ‘marine’ or ‘freshwater’ based on their collection location). To account for non-independence of observations within studies as well as methodological variation among studies, we included study ID as a random effect on the intercept in all models. To examine additional variation in *T*_*opt*_ across performance types after accounting for latitudinal patterns, realm, and study-level differences, we analyzed residuals from the latitude-*T*_*op*t_ model (hereby referred to as ‘residual thermal optimum’).

To assess whether activation energy patterns were consistent with previous work, we restricted analyses to datasets with sufficient information on the rising limb of the thermal performance curve. Specifically, we included curves classified as full curve, optimum-only, full-rise-with-optimum, full-rise-without-optimum, or partial-rise. We fit a Boltzmann-Arrhenius model to the rising portion of each thermal performance curve (raw data at temperatures ≤ *T*_*opt*_ ) using nonlinear least squares. Curves with fewer than four observations on the rising limb were excluded to ensure sufficient data for stable estimation of arrhenius parameters and to reduce unreliable model fits. Fits with r^2^ > 0.5 and P < 0.05 were considered as sufficiently well-fit and retained for subsequent analysis [16].

In order to understand differences in thermal responses depending on the context of the ecological performance response, we compared residual thermal optima and activation energy across grouping schemes: response type, motivation, and level of biological organization. We evaluated group differences using separate one-way generalized linear models. We did not expect a pattern between absolute latitude and activation energy, and thus did not analyze latitude-corrected activation energy (as was done for thermal optima; see results for confirmation).

We also fit linear mixed effect models to residual thermal optimum and activation energy to test whether (a) thermal optima increase with thermal breadth under the “broader is hotter” hypothesis [28], and (b) activation energy varies systematically with thermal breadth under “stenothermal is steeper” hypothesis [30]. We held study ID as a random effect in each model, and considered separately the fixed predictors of performance breadth and thermal tolerance.

## Results

### Data characteristics

We assembled a large and geographically diverse dataset of thermal-performance responses and there were often multiple curves per species (Figure 2C and D). We extracted 457 thermal performances datasets from 118 different studies (1978-2024), representing 107 fish species from 138 collection locations (Figure 3). While capturing a wide geographic range, most sampling occurred coastally and across North America and Western Europe, with relatively little data from tropical species. Fish from marine habitats are most represented in the data (254 datasets; includes coastal, oceanic, or intertidal habitats), followed by fish coming from freshwater (185 datasets; includes river, lake, or pond habitats) and brackish (18 datasets; includes estuaries, mangroves, or lagoon habitats) environments. Although studies on marine and freshwater were roughly equal in number (60 and 58, respectively), the number of datasets differs because studies contribute uneven amounts of data. Some studies measured several performance traits (e.g. swimming speed and metabolic metrics) within the same study. Others evaluated the same trait across multiple treatments, or measured performance across multiple species. As a result, individual studies often yielded multiple thermal-performance responses. On average, each study contributed 3.87 datasets (range: 1-20), meaning that a relatively small number of studies produced a disproportionately large share of the total data.

Our data represents considerable variation in species, response, and experimental paradigm. The compiled data span species from 43 fish families, with Gadidae (Cod), Lutjanidae (Snapper), Salmonidae (Salmonid), and Leuciscidae (Minnow) families as the four most commonly studied (Figure S1). Across studies, the most frequently measured physiological responses were metabolism (141), somatic growth (122), locomotion (89, Figure 3B). When categorized by underlying trait motivation, metabolic traits encompassed autonomic (eg. resting metabolism), voluntary (aerobic scope), and negative (maximum metabolic rate) responses. In contrast, somatic growth traits were all autonomic, baseline physiological responses. Locomotion traits were most often negatively motivated, commonly measured through burst swimming speed elicited by a startle or escape response (Figure 3B).

Most data were collected from juveniles (214 datasets) and adults (104). Early life stages (embryos, larvae, fry) were less frequently tested, making up fewer than 10 % of the compiled data. Over half of the datasets (253) involved no experimental manipulation beyond temperature. Among datasets with additional treatments (204), food ration (51), salinity (45), and acclimation temperature (43) were the most common.

Studies differed in the time of exposure to test temperatures and level of reporting. Most (323) were batch-acclimated, in which groups of fish were reared at the same temperature that performance was measured at (e.g. growth trials conducted in separate 10℃, 15℃, 20℃, 30℃ tanks). The remaining datasets measured acute thermal responses, where individuals were exposed to multiple temperatures in ramping or short-duration trials. Finally, most datasets provided mean (414) or median (3) performance values at each temperature, whereas a smaller subset reported individual-level measurements (40).

### Curve fitting & coverage

The model with the lowest AIC varied among the datasets. Among curves with four test temperatures, the Quadratic model [47] was often the best fit (134) compared to the Gaussian (65) [48]. The most common model fit for datasets with above four test temperatures varied more, but Spain (51) [49] and Weibull (50) [11] were most frequent, followed closely by several other four-parameter models (Figure S2).

We found substantial variation among studies in the number and range of test temperatures used, leading to differences in how much of the thermal response was observed in each assay. The median number of unique temperatures tested was 4 (this was also the minimum), and the mean was 5.1, with only a quarter of the datasets (26.5 %) sampled at more than five temperatures (Figure S1). From this, we found most datasets covered sufficient aspects of the thermal performance curve to identify a single local maximum (*T*_*opt*_ ), and/or to estimate the rate of performance increase with temperature (Figure 2B). Only datasets with more than 5 temperatures were among those that met criteria of a full curve, i.e. with observations of performance values at or below 20% of maximum performance values on either side of an optimum (Figure 2B). This yielded fewer estimates of thermal tolerance breadth (defined as the temperature range over which performance remained at or above 10 % of the maximum rate, n=26) compared to thermal performance breadths (temperature range over which performance remained at or above 80 % of the maximum rate, n = 96), but these parameters were highly correlated among the fish for which we had both (Figure S3). The number and distribution of non-pseudoreplicated thermal performance curves (i.e. the average curve for each response type, species, and latitude) was similar in marine vs. freshwater realms, and shifted to cooler temperatures with increasing latitude (Figure 2C and D).

### Relationships between curve parameters, latitude, and experienced temperatures

We detected several expected relationships between curve parameters, latitude, and experienced temperatures. Thermal optima declined with latitude and increased with mean annual temperature, and did so with a steeper slope in marine compared to freshwater fish (slope_marine_ = -0.42 ± 0.06 and 0.87 ± 0.13; slope_freshwater_ = -0.18 ± 0.04, and 0.46 ± 0.11, Figure 4A-B, Table S1). We also found thermal optima increased with upper environmental temperatures significantly in marine but not freshwater fish (slope_marine_ = 0.83 ± 0.14, and slope_freshwater_ = 0.17 ± 0.11, Figure S4). Thermal performance breadth, defined as the temperature range over which performance remained at or above 80 % of the maximum rate, increased with temperature variation in freshwater but not marine habitats (slope_freshwater_ = 0.70, slope_marine_ = 0.34, Figure 4D, Table S1), although the pattern with latitude, if present, was underpowered (Figure S5). We did not find support for a relationship between Thermal Safety Margin (TSM) and temperature variation (Figure 4E, Table S1), although we did find that TSM is greater altogether among freshwater fish compared to marine (Figure 4E), and increases with latitude among freshwater fish (slope = 0.14 ± 0.04, Figure 4C, Table S1).

**Figure 4.**
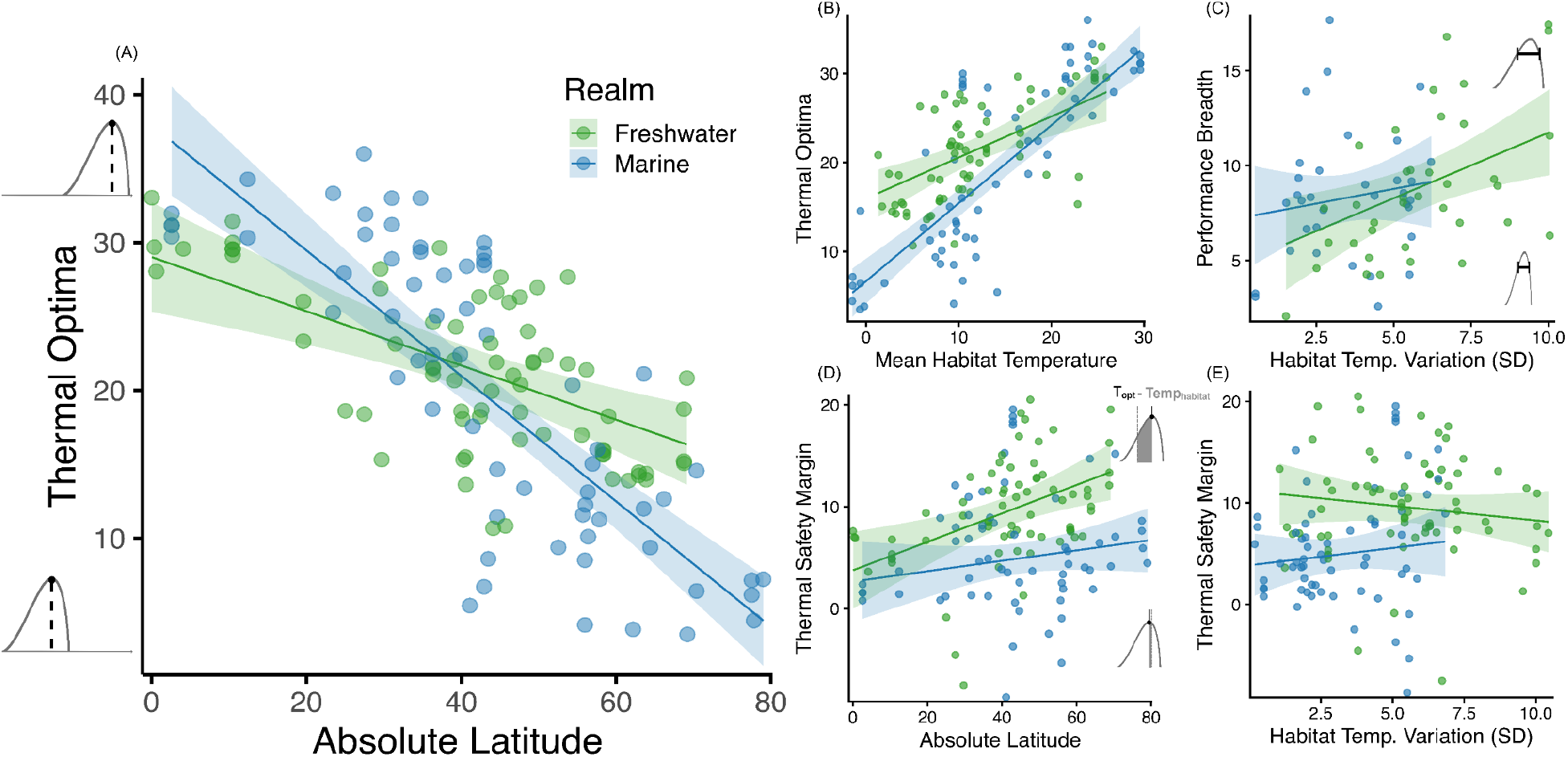
Environmental predictors of thermal trait variation across semi aggregated thermal performance curves. (A). Relationship between thermal optimum and latitude. (B). Relationship between thermal optima and environmental water temperature at the collection location, quantified using mean monthly temperature from 1982-2025. (C). Relationship between performance breadth (temperatures supporting above 80% of maximum performance) and thermal variability, measured as the standard deviation (SD) of mean monthly temperatures. (D). Relationship between thermal safety margin (TSM = *T*_*opt*_ - *T*_*habitat*_) and absolute latitude, where *T*_*habitat*_ is the mean monthly temperature at the collection location. (E) Relationship between thermal safety margin and thermal variability. Lines show fitted relationships from linear mixed-effects models. Shaded bands represent 95% confidence intervals, and points represent raw data, colored by realm, freshwater (green) and marine (blue). Small curve schematics alongside the y-axes in panels (A), (C), and (D) illustrate thermal traits on a TPC.

After accounting for absolute latitude, realm, and study-level effects, residual thermal optima differed across performance classifications. We found that metabolic responses had thermal optima that were 3.48± 0.94 times greater than those of reproductive responses (Figure 5A, Table S2). Thermal optima differed across levels of biological organization, with population-level responses showing lower thermal optima compared to internal responses (holding latitude and realm constant, by an effect size of -2.84 ± 0.79, Figure 5A, Table S2). Residual thermal optima were also lower among fish with narrower performance and tolerance breadths (Figure S6). We did not have an expectation that residual thermal optima differed among motivation categories and did not find evidence for any differences of note.

**Figure 5.**
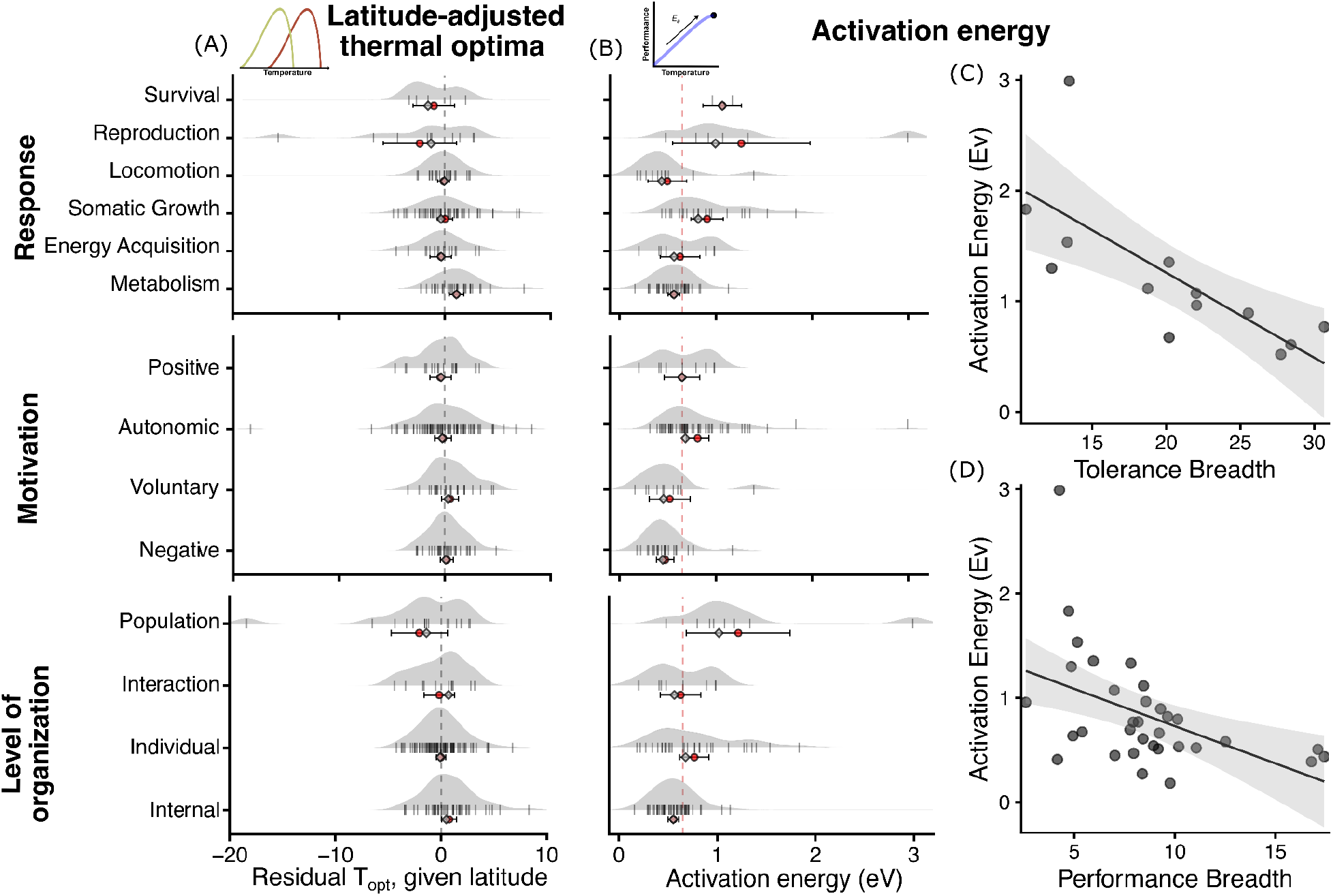
Latitude-adjusted thermal optima and activation energy across performance classifications. (A) Residual thermal optima after accounting for absolute latitude, realm (marine vs. terrestrial), and study-level variation in a linear mixed effects model (give model). Grey ridgelines, rug marks, and summary statistics are shown as in panel A. The grey dashed vertical line is at x = 0 variation. (B) Distributions of activation energy estimates (Ea, eV) for datasets exhibiting an eligible rise in performance grouped by trait physiology, motivation, and level. Grey ridgelines show density dataset-level estimates, with rug marks indicating individual observations. Red points denote group means, and light grey points denote medians, with associated uncertainty (±95%). The red dashed vertical line indicates the MTE expectation (0.65 eV). (C) Relationship between thermal tolerance breadth (*T*_*min*_ to *T*_*max*_ range) and activation energy. Slope = -0.08 ± 0.02 (Table S4). (D) Relationship between thermal performance breadth (temperatures supporting above 80% of maximum performance) and activation energy. Slope = -0.07 ± 0.02 (Table S4).

The positioning of other curve parameters, such as the temperature range of increasing and decreasing portions of the curves, the temperature of performance breadth maxima and minima (i.e. pejus temperatures), and maximum and minimum endpoints of positive performance (thermal tolerance minima and maxima), each showed patterns with latitude consistent with adaptation to thermal variation, but with fewer data with which to test for patterns (Figure 6).

**Figure 6.**
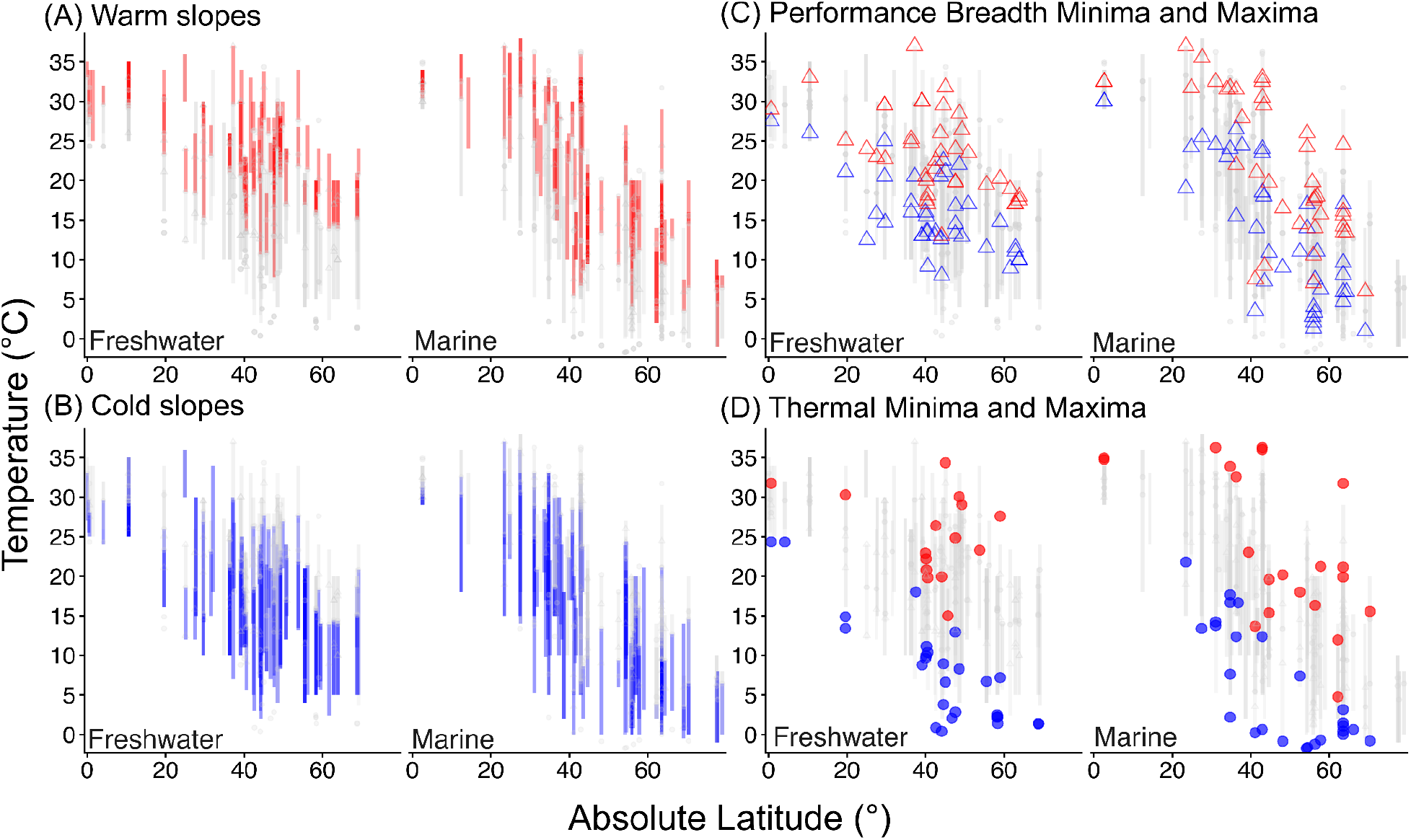
Latitudinal variation in thermal performance curve parameters across freshwater and marine fishes. (A) ‘Declining side’ of thermal responses plotted against absolute latitude. Lines represent assayed portions of the thermal response that are warmer than the optimal temperature or in which responses exhibit monotonic decline. (B) ‘Rising side’ thermal responses. Lines represent assayed portions of the thermal response that are cooler than the optimal temperature or responses exhibiting monotonic rise. (C) Performance breadth minima (blue triangles) and maxima (red triangles) representing temperatures at which performance declines to 80% of the maximum performance at Topt. An upper and lower estimate along the same curve (red and blue triangles on the same grey line) display the estimated performance breadth. (D) Thermal tolerance minima (blue circles) and maxima (red circles) across latitude. Greyed elements in all panels indicate other curve parameters along assay temperatures to help illustrate connections.

### Activation Energy of Performance Rises

The median activation energy of rises in performance was 0.65 eV, but the mean was 0.71 ± 0.05 eV indicating a right-skew. There was also substantial variability. When comparing among categories of performance types, metabolic responses had a mean activation energy of 0.56 ± 0.03eV (median = 0.56 eV) that was closest to the central tendency predicted by MTE (0.65eV for MTE, [14]). However, other performance types had either lower or higher activation energies (Figure 5A, Table S3). We found that negatively motivated traits exhibited significantly lower activation energies than autonomic traits (Figure 5B, Table S3). Similarly, internal traits showed significantly lower activation energy than those operating at the individual or population level (Table S3), although the interaction-level activation energy was not different from either (Table S3). Within most groupings, the median value was lower than the mean value, indicating a right skew in the distribution (Figure 5A), although our models were fit with a gamma distribution for positive non-zero values thus accounting for skew in residual variation. Activation energy increased significantly with performance breadth (range of temperatures that span 80% of maximum performance) and with tolerance breadth (range of temperatures that span 10% of maximum performance; Figure 5C-D, Table S4), but not with latitude (Figure S7).

## Discussion

The FishTherm dataset provides a comprehensive aggregation of thermal performance assessments of freshwater and marine fishes. Parsing the 457 datasets by the portions of thermal responses from which comparable parameters can be estimated yields an accessible and comparative trait dataset for testing hypotheses in the evolutionary ecology of thermal performance while characterizing a valuable response trait in fish. Our finding that fish thermal optima increase towards the equator and in warmer waters is consistent with expectations of thermal adaptation and previous findings in other systems [17,18,35], and by accounting for latitude we find new support for the hypothesis that thermal optima are greater in responses measured at lower levels of biological organization [26,27,50]. Additionally, we find systematic variation in activation energies among response types and curve shapes that provide new evidence for the life-dinner principle [16], the complexity-reducing-TPC hypothesis [27], and a positive relationship between thermal performance breadth and temperature sensitivity that is consistent with a Generalist-Specialist trade off hypothesis [28,30]. Here we expand on our findings in relation to these hypotheses, discuss how our results contribute new insights into geographic variation in thermal responses, and highlight important caveats and opportunities for further analysis of this new dataset.

Our findings suggest that marine fish have more variable thermal optima that track latitude and mean environmental temperature more closely compared to freshwater species. While less variable, the thermal optima of freshwater fishes are almost always higher than the mean environmental temperature at each collection location, leading to a thermal safety margin (*T*_*opt*_ - *T*_*habitat*_) that is greater on average than in marine species, and increases with latitude. This suggests that freshwater species’ thermal optima do not conform to environmental temperatures as closely as marine species, but remain well above mean annual temperature. We had expected freshwater habitats to have greater thermal safety margins due to higher environmental variability [18], but also expected thermal safety margins to increase with environmental variability within each realm, which we did not find. Instead, this freshwater-marine difference in the positioning of *T*_*opt*_ relative to mean annual temperature could reflect the more seasonal activity patterns expected in freshwater systems. Rather than adapting to an annual mean, the thermal optima of freshwater fishes at higher latitudes could represent adaptation to summer (active) temperatures, hence they are altogether higher. Another explanation could be a greater level of spatial heterogeneity expected in freshwater systems. Through microhabitat selection, freshwater fishes might maintain higher body temperatures and therefore maintain higher and less variable *T*_*opt*_ (the Bogert effect [22]). In addition, we cannot rule out the possible effects of different sources of environmental temperature estimated for marine vs. freshwater realms. We used satellite-derived measures of sea surface temperature for marine species, and temperatures derived from a hydrological streamflow model for freshwater fish; these different modelling approaches could lead to systematically different mean temperatures, and potentially different estimates of variation. Still, our finding of broader thermal performance breadths in more temporally variable freshwater environments is consistent with expectations under the climate variability hypothesis [51] and with previous results in fish [30], and suggests thermal breadths but not thermal safety margins respond to variable temperatures in freshwater fish.

Our results provide new evidence for the hypothesis that responses at higher orders of biological complexity have lower thermal optima and steeper sensitivities towards those optima. It has long been expected that thermal tolerance ranges for growth, activity, and reproduction become increasingly narrow at higher levels of organization, in part because of the longer duration or loading of stress accrued by higher-level processes [52], and in part because more complex systems are limited by more concurrent processes [29]. Our results are consistent with a previous synthesis of thermal performance curves in ectotherms that found higher-level processes to have a higher exponential rise in performance towards a thermal optimum (i.e. activation energy), although this was mainly supported in insects and reptiles, with a paucity of data at the time for fish [16]. A compilation of ‘full’ thermal performance curves in insects, reptiles, and plants provided more evidence that thermal performance curves were narrower and cold-shifted at higher levels of organization, although thermal performance curves for only two levels of organization were available within each group at that time [26]. By experimentally assessing three levels of biological organization in a single insect (*Drosophila melanogaster*), Bozinovic et al. showed that TPCs were more narrow and had a cooler *T*_*opt*_ with increasing level of organization and proposed a mechanistic framework by which biological rates at higher levels emerge from the interaction of lower level responses that are both variable and skewed [27]. Our results extend this work by providing new comparative analyses of activation energies, thermal optima, and thermal breadths that span four levels of biological organization in fish. Our finding that population-level responses, such as survival and reproduction, increase at a faster rate towards a colder optimum as compared to individual and internal processes is a new finding in support of this hypothesis.

Our findings provide new insights into the constraints and sources of variability of the fundamental metabolic activation energy scaling constant. We show that activation energies in fish vary in systematic ways away from the empirical central tendency of 0.65eV [14], and in line with directions predicted and observed in previous studies. As predicted under the life-dinner principle [24,16], we found that negatively-motivated responses such as escape performance have a reduced slope compared to autonomic responses, such as basal metabolic rate. This adds to evidence compiled from insects, lizards, mammals and fish [16], observable only by scoring response curves according to motivation type following the ontogeny of Dell et al. Our results also support the idea that activation energy is constrained by TPC breadth. Consistent with Dahlke et al., who reported this pattern for developmental rates relative to critical thermal limits in fishes [30], we found that responses with wider thermal breadths also exhibited steeper activation energy, across a broader range of response types. We believe these results suggest limits to the conservation of activation energy potentially driven by a trade-off between thermal breadth and maximum performance [53], although we were not able to compare the heights of TPCs in our dataset directly.

By fitting models to categorized data types and carefully assessing thermal performance curves, our analyses provide insights about variation in thermal performance across fish. All data included in model fitting met strict criteria and passed further visual inspections. Still, even among the datasets with enough information to build full thermal performance curves, we found that among 11 TPC models considered, not one model fit every dataset, nor were there obvious patterns that connected model types to particular performance types. We had examples of responses that were right-skewed rather than left-skewed, matching variation also reported in phytoplankton curve shapes [54]. The lack of a single best model is in line with Kontopoulos et al. [37] and suggests that TPCs of fish do not fit a universally-shaped thermal performance curve [55]. Although only 6% of our datasets were sufficient for estimating a full TPC, we were able to leverage many more datasets, including performance curves with as low as 4 temperatures, to estimate common curve parameters (*T*_*opt*_, eV), by mapping datasets to other components of the TPC. Future work could focus on further investigating the variation in curve fits and variation (or uniformity) in curve shape across responses and trying to parse apart the drivers of such variation.

Beyond the patterns we present, FishTherm includes additional characteristics of interest for future analysis. We extracted and codified metrics about acclimation conditions as well as duration of treatments whenever available. This included experiments repeated at different acclimation temperatures, which could be leveraged to further refine comparative analyses and test further hypotheses about acclimation responses [56] and the impacts of temporal dynamics on TPCs [57]. We also extracted and codified other environmental conditions that varied either within or across studies, such as oxygen level and salinity, which could be leveraged to further understand responses to interacting stressors [58,59]. Similarly, we noted information about body size and life-history stages used in each treatment, for the purpose of eventually testing how these factors affect TPC shape [30], and recorded values of sample size and estimates of precision (e.g. standard error) to allow precision-based weighting factors for more refined analyses [42]. We did not attempt to transform response data into equal units, but unit conversion with similar response types could allow curve heights to be more directly compared [26,53].

Our results present informative and potentially consequential patterns of variation in thermal responses of fishes across scales of organization. Narrower and more sensitive thermal performance curves at increasing biological scales means that when we extrapolate results from laboratory experiments to the scales where nature conservation and management typically happen (e.g., population, community), we likely overestimate thermal performance at the warm end of the curve and underestimate sensitivities (slope of rise) at the cold end of the curve. We hope this dataset can lead to many further discoveries and applications, particularly given the widespread changes in fish distributions globally and their importance to global food security.

