## Supplemental Materials for "Thermal performance in fishes varies systematically across latitude, habitat, and biological organization"

Table of contents

*Supplementary Methods*……………………………………………………………………………………………..
List of search terms ……………………………………………………………………………………..2

*Supplementary Figures…*…..…………………………………………………………………………
Figure S1 - Taxonomic representation and temperature treatment structure across compiled data……..….3

#### Supplementary Methods

##### List of search terms

Search Terms used in our literature search for subsequent systematic review were as follows:

1. **Pilot search terms:**
   TS=((fish* OR teleost*) AND (therm* OR temperature*) AND (performance* OR breadth OR window* OR optim* OR "performance curve*" OR biologg* OR accelerometer* OR "swim* speed*" OR "swim* velocity" OR growth OR "growth rate*" OR "population growth" OR fitness OR "reproductive success" OR gonad*))

2. **Final search terms:**
   (TS=(fish OR teleost)) AND TS=("thermal response" OR "temperature response" OR "thermal range" OR "thermal tolerance" OR "thermal performance" OR temperature NEAR/6 effect OR temperature NEAR/6 influence OR temperature NEAR/6 relation OR "water temperatures") AND TS=(performance OR swim* OR speed OR velocity OR reproduction OR reproductive OR growth OR activity OR behavio$r OR feeding OR prey OR escape))

#### Supplementary Figures

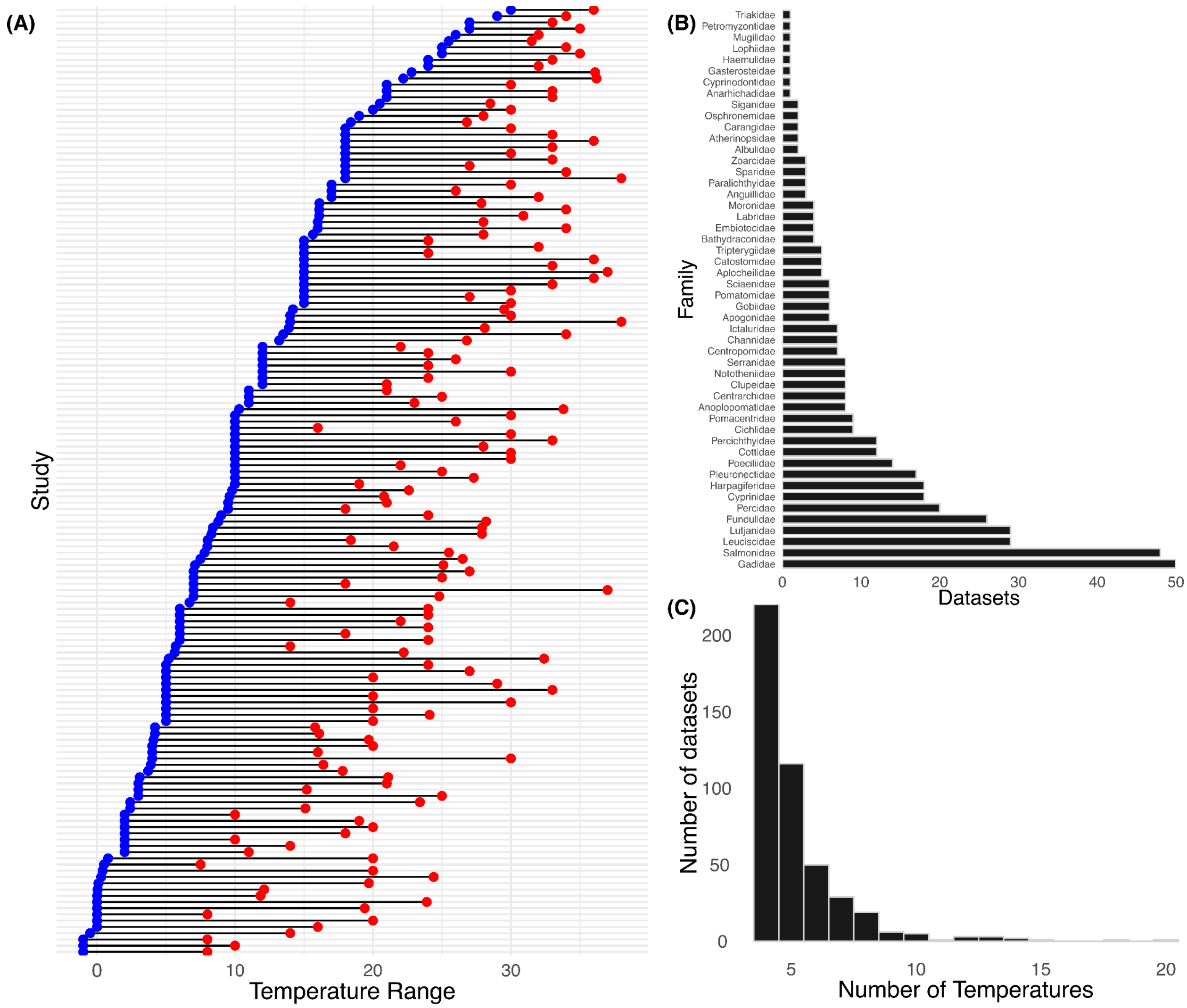

Figure S1. Taxonomic representation and temperature treatment structure across compiled thermal performance curve data. (A) Minimum (blue) and maximum (red) experimental testing temperatures are linked by black lines showing the range tested per study. Studies are ordered by minimum temperature. (B) Number of thermal performance curve datasets represented per fish family across the full database (n = 457). (C) Frequency distribution of the number of unique temperature treatments per dataset.

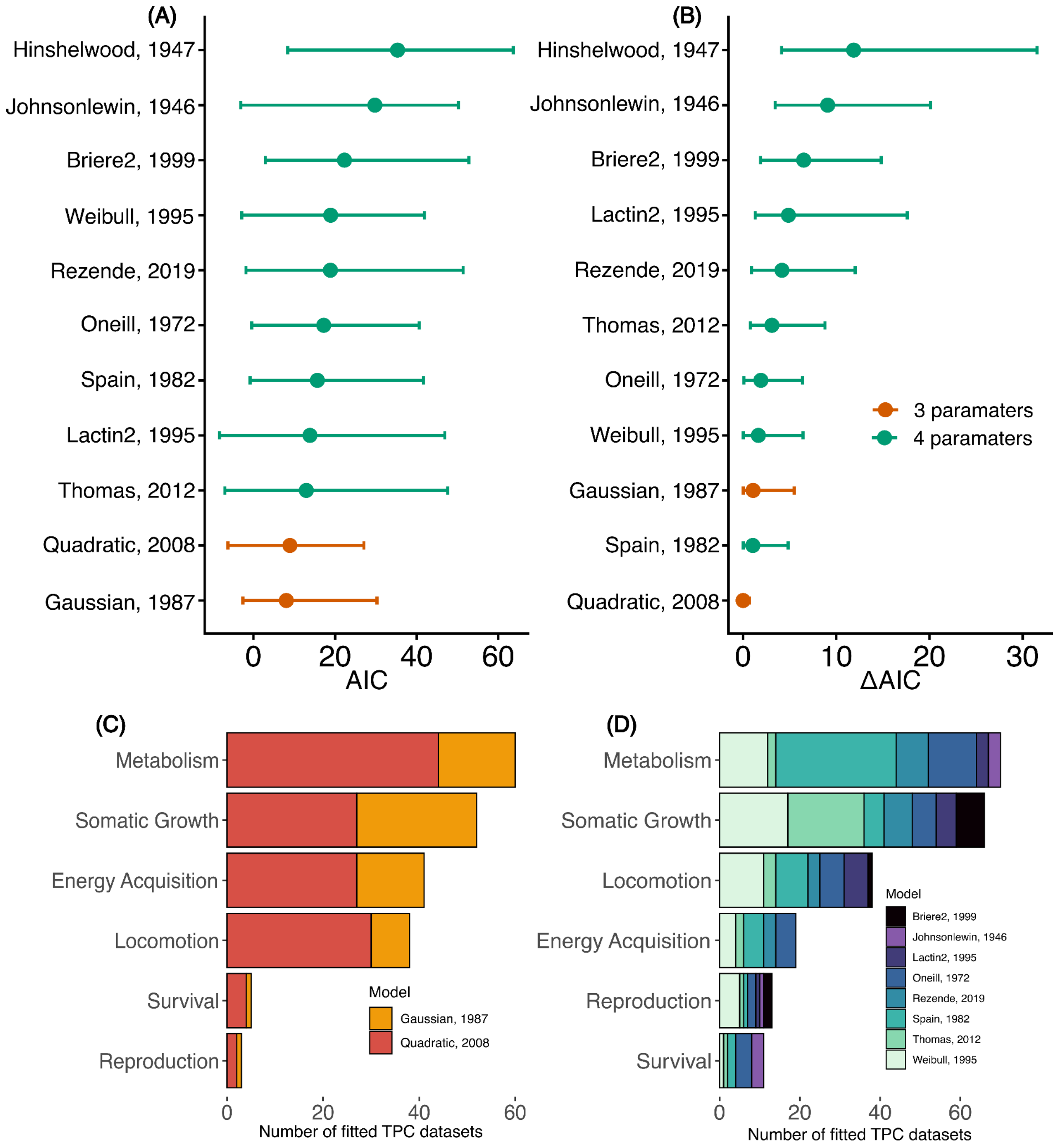

Figure S2. Candidate model performance and best-fit model across compiled data. Mean AIC values and 95 % CI for all candidate thermal performance curve models evaluated, with colors indicating three-parameter (orange) and four-parameter (turquoise) models. (B) Mean ΔAIC values for all candidate models, calculated as the difference between each model's AIC and the lowest AIC within each dataset, with colors indicating three-parameter (orange) and four-parameter (turquoise) models. (C) Best-fitting three-parameter model (lowest AIC) for each low-resolution dataset, grouped by response type and colored by model identity. (D) Best-fitting four-parameter model (lowest AIC) for each high-resolution dataset, grouped by response type and colored by model identity.

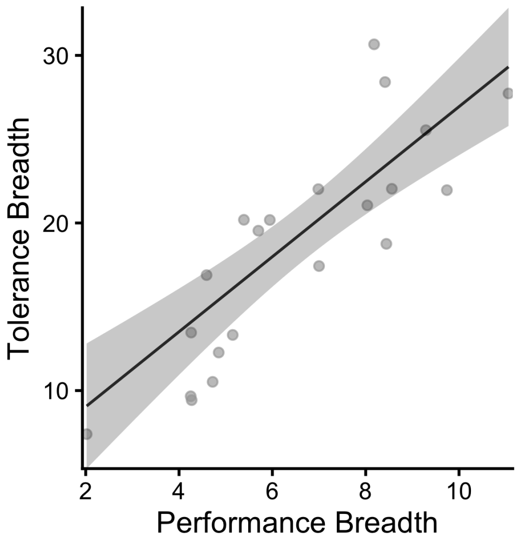

Figure S3. Relationship between performance breadth and tolerance breadth. Line and associated 95% confidence interval represent predictions from a linear mixed model effects model with study ID included as a random intercept, while points represent raw data from aggregated curves. Tolerance breadth increased significantly with performance breadth (slope = 2.24 ± 0.36 SE, $p = 6.3 \times 10^{-6}$).

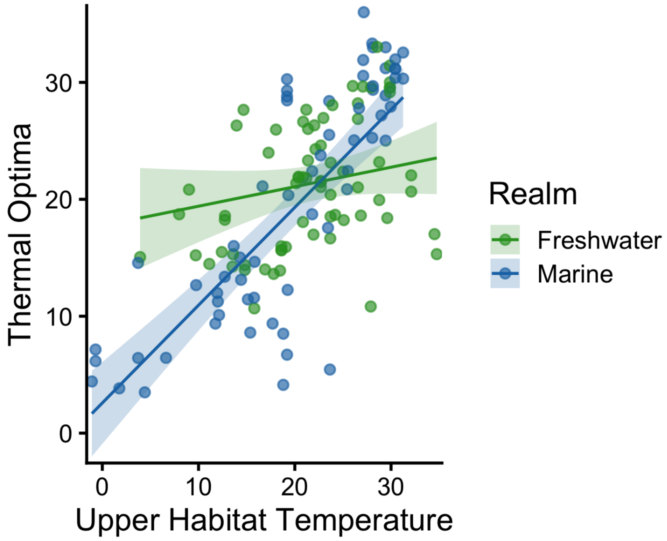

Figure S4. Relationship between upper habitat temperature and thermal optima. Lines and associated 95% confidence intervals represent predictions of $T_{opt}$ from a linear mixed-effects model including latitude, environment (marine vs freshwater), and their interaction, with study ID included as a random intercept (n = 134 observations, 88 studies), while points represent raw data aggregated curves. $T_{opt}$ did not significantly increase with upper habitat temperature in freshwater systems (slope = 0.17 ± 0.11 SE, p = 0.135) but did in marine systems (slope = 0.83 ± 0.14, p = 4.04 × $10^{-6}$).

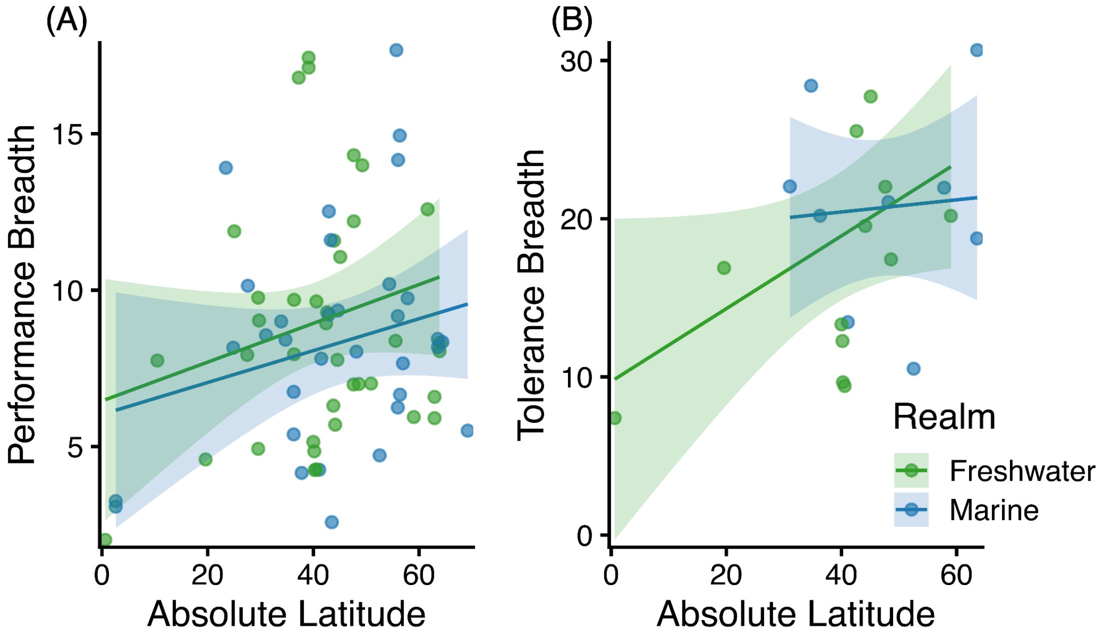

Figure S5. Relationship between absolute latitude and (A) performance and (B) tolerance breadth. (A). Lines and associated 95% confidence intervals represent (A) predictions of performance breadth (80% maximum performance) and (B) predictions of tolerance breadth from linear mixed-effects models including latitude, environment (marine vs freshwater), and their interaction, with study ID included as a random intercept (performance breath: n = 71 observations, 54 studies; tolerance breadth: n = 21 observations, 17 studies) . Points represent raw data from aggregated curves. No significant relationship between latitude and performance breadth or tolerance breadth was detected in freshwater or marine systems (performance breadth: $slope_{freshwater} = 0.06 \pm 0.05$ SE, $p_{freshwater} = 0.19$, $slope_{marine} = 0.05 \pm 0.06$ SE, $p_{marine} = 0.86$; tolerance breadth: $slope_{freshwater} = 0.23 \pm 0.12$ SE, $p_{freshwater} = 0.09$, $slope_{marine} = 0.04 \pm 0.19$ SE, $p_{marine} = 0.33$) but sample sizes were limited for both of these analyses, particularly at low latitudes.

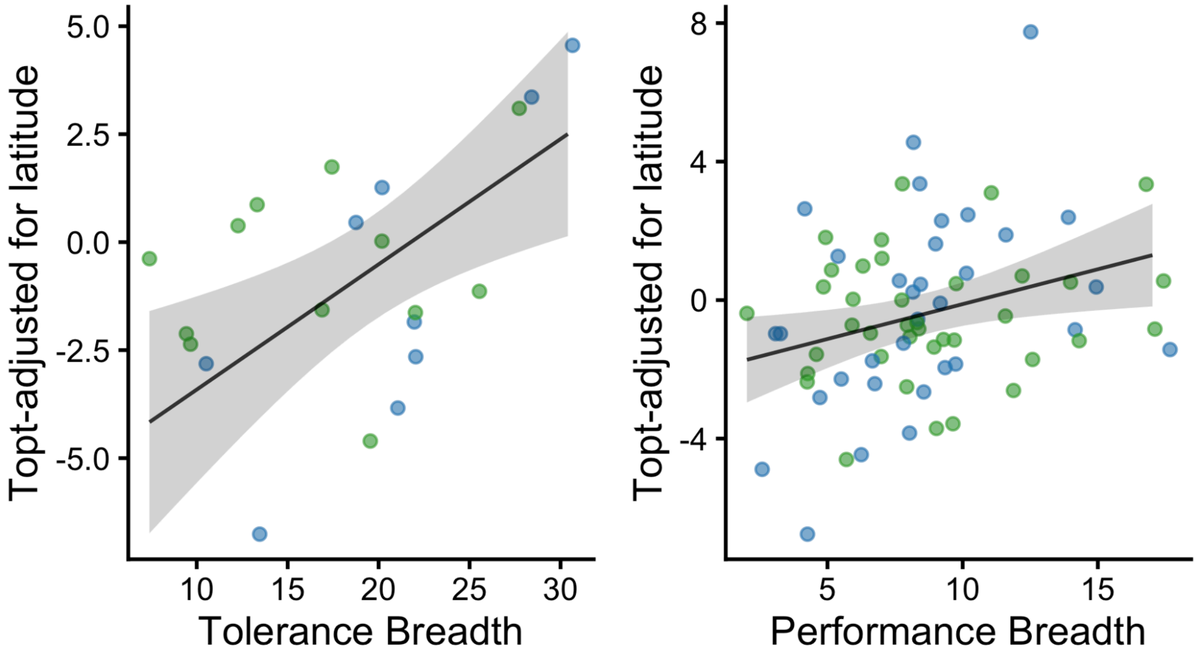

Figure S6. Relationship between (a) thermal tolerance breadth and latitude-adjusted thermal optima ($T_{opt}$), and (b) performance breadth and latitude-adjusted $T_{opt}$. Latitude-adjusted $T_{opt}$ values represent residual variation after accounting for absolute latitude in a mixed-effects model (with study ID included as a random effect). Points represent aggregated TPCs, colored by realm: freshwater (green) and marine (blue); solid lines show linear model fits with shaded areas indicating 95% confidence intervals. Thermal tolerance breadth was positively associated with latitude-adjusted $T_{opt}$ (slope = 0.29 ± 0.10 SE, p = 0.007), and performance breadth also showed a positive relationship (slope = 0.20 ± 0.08 SE, p = 0.018).

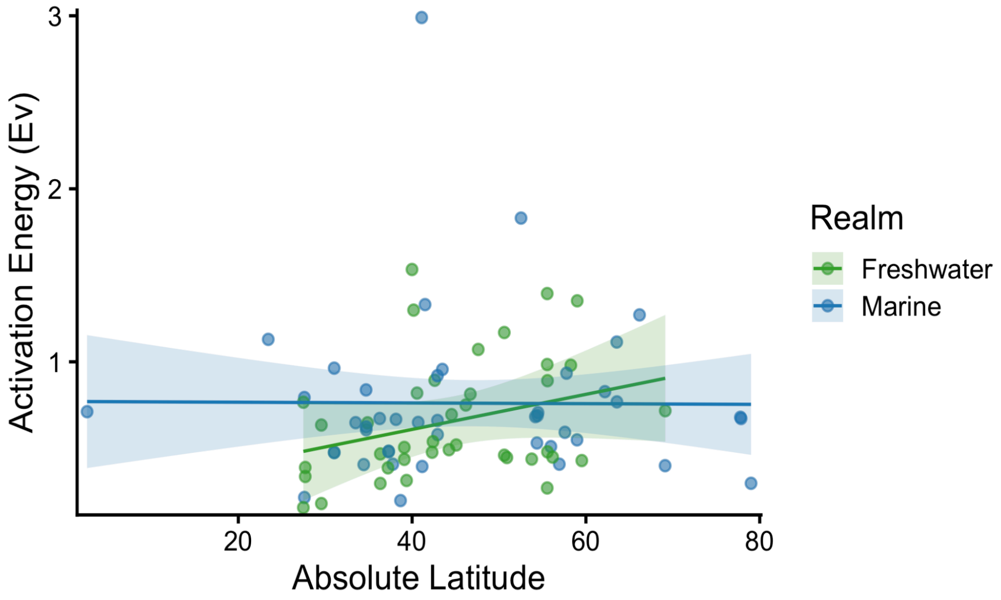

Figure S7. Relationship between activation energy and absolute latitude. Lines and associated 95% confidence intervals represent predictions from a linear mixed-effects model including latitude, environment (marine vs freshwater), and their interaction, with study ID included as a random intercept (n = 83 observations, 64 studies), while points represent raw data. Activation energy did not vary significantly with latitude in either freshwater (slope = 0.010 ± 0.007 SE, p = 0.164) or marine systems (slope = −0.00021 ± 0.008 SE, p = 0.216), and the latitude × environment interaction was not significant.

### Supplementary Tables

| | $T_{opt}$ | | | | | $T_{opt}$ | | | | | Thermal Safety Margin | | | | | Thermal Safety Margin | | | | | Performance Breadth | | | | |
|---|---|---|---|---|---|---|---|---|---|---|---|---|---|---|---|---|---|---|---|---|---|---|---|---|---|
| *Predictors* | *Estimates* | *std. Error* | *CI* | *Statistic* | *p* | *Estimates* | *std. Error* | *CI* | *Statistic* | *p* | *Estimates* | *std. Error* | *CI* | *Statistic* | *p* | *Estimates* | *std. Error* | *CI* | *Statistic* | *p* | *Estimates* | *std. Error* | *CI* | *Statistic* | *p* |
| (Intercept) [Freshwater] | 29.01 | 1.90 | 25.25 – 32.77 | 15.26 | **<0.001** | 15.95 | 1.49 | 13.01 – 18.89 | 10.73 | **<0.001** | 3.76 | 1.92 | -0.05 – 7.56 | 1.95 | 0.053 | 11.17 | 1.81 | 7.58 – 14.75 | 6.17 | **<0.001** | 4.77 | 1.59 | 1.60 – 7.94 | 3.00 | **0.004** |
| Abs latitude x Freshwater | -0.18 | 0.04 | -0.27 – -0.10 | -4.30 | **<0.001** | | | | | | 0.14 | 0.04 | 0.05 – 0.23 | 3.23 | **0.002** | | | | | | | | | | |
| (Intercept) [Marine] | 8.93 | 2.76 | 3.46 – 14.40 | 3.23 | **0.002** | -9.26 | 1.97 | -13.16 – -5.37 | -4.70 | **<0.001** | -1.09 | 2.80 | -6.63 – 4.44 | -0.39 | 0.663 | -7.33 | 2.44 | -12.16 – -2.50 | -3.00 | **0.003** | 2.38 | 2.19 | -2.00 – 6.75 | 1.09 | 0.282 |
| Abs latitude x Marine | -0.24 | 0.06 | -0.36 – -0.12 | -4.08 | **<0.001** | | | | | | -0.09 | 0.06 | -0.21 – 0.03 | -1.48 | 0.141 | | | | | | | | | | |
| Mean Temp x Freshwater | | | | | | 0.46 | 0.11 | 0.24 – 0.68 | 4.20 | **<0.001** | | | | | | | | | | | | | | | |
| Mean Temp x Marine | | | | | | 0.41 | 0.13 | 0.15 – 0.68 | 3.08 | **0.003** | | | | | | | | | | | | | | | |
| Thermal Variation (SD) x Freshwater | | | | | | | | | | | | | | | | -0.29 | 0.29 | -0.85 – 0.27 | -1.01 | 0.312 | 0.70 | 0.24 | 0.21 – 1.19 | 2.86 | **0.006** |
| Thermal Variation (SD) x Marine | | | | | | | | | | | | | | | | 0.64 | 0.52 | -0.40 – 1.67 | 1.21 | 0.227 | -0.36 | 0.46 | -1.28 – 0.56 | -0.78 | 0.436 |
| **Random Effects** | | | | | | | | | | | | | | | | | | | | | | | | | |
| $\sigma^2$ | 11.94 | | | | | 12.00 | | | | | 13.02 | | | | | 13.89 | | | | | 12.17 | | | | |
| $\tau_{00}$ | 14.82 $_{study\_ID}$ | | | | | 12.83 $_{study\_ID}$ | | | | | 14.56 $_{study\_ID}$ | | | | | 15.98 $_{study\_ID}$ | | | | | 8.91 $_{study\_ID}$ | | | | |
| ICC | 0.55 | | | | | 0.52 | | | | | 0.53 | | | | | 0.54 | | | | | 0.80 | | | | |
| N | 88 $_{study\_ID}$ | | | | | 88 $_{study\_ID}$ | | | | | 88 $_{study\_ID}$ | | | | | 88 $_{study\_ID}$ | | | | | 54 $_{study\_ID}$ | | | | |
| Observations | 134 | | | | | 134 | | | | | 134 | | | | | 134 | | | | | 71 | | | | |
| Marginal $R^2$ / Conditional $R^2$ | 0.580 / 0.813 | | | | | 0.583 / 0.799 | | | | | 0.250 / 0.646 | | | | | 0.162 / 0.610 | | | | | 0.116 / 0.827 | | | | |

Table S1. Mixed-effects model results for models fit to thermal performance curve parameters as a function of latitude and environmental temperatures in freshwater and marine fish. Linear mixed-effects models were fit separately to $T_{opt}$, thermal safety margin, and performance breadth. Fixed effects included absolute latitude, mean environmental temperature, and thermal variation (SD), each interacted with the environment (freshwater vs marine). Study ID was included as a random effect on the intercept. Reported are fixed-effect estimates with standard errors, 95% confidence intervals, test statistics, and p-values. Marginal and conditional $R^2$ values are shown for each model.

| | *Response Type* | | | | | | *Response Motivation* | | | | | | *Response Organization* | | | | |
|---|---|---|---|---|---|---|---|---|---|---|---|---|---|---|---|---|---|
| | **resid $t_{opt}$** | | | | | | **resid $t_{opt}$** | | | | | | **resid $t_{opt}$** | | | | |
| *Predictors* | *Estimates* | *std. Error* | *CI* | *Statistic* | *p* | | *Estimates* | *std. Error* | *CI* | *Statistic* | *p* | | *Estimates* | *std. Error* | *CI* | *Statistic* | *p* |
| Intercept (Metabolism) | 1.13 | 0.46 | 0.22 – 2.04 | 2.44 | **0.015** | Intercept (Negative) | 0.22 | 0.51 | -0.77 – 1.21 | 0.44 | 0.663 | Intercept (Internal) | 0.76 | 0.40 | -0.02 – 1.55 | 1.91 | 0.056 |
| Energy aquisition | -1.50 | 0.79 | -3.05 – 0.06 | -1.89 | 0.059 | | | | | | | | | | | | |
| Somatic growth | -1.10 | 0.59 | -2.26 – 0.06 | -1.86 | 0.064 | | | | | | | | | | | | |
| Locomotion | -1.23 | 0.71 | -2.62 – 0.16 | -1.73 | 0.084 | | | | | | | | | | | | |
| Reproduction | -3.48 | 0.94 | -5.32 – -1.64 | -3.72 | **<0.001** | | | | | | | | | | | | |
| Survival | -2.14 | 1.24 | -4.58 – 0.29 | -1.73 | 0.084 | | | | | | | | | | | | |
| Voluntary | | | | | | | 0.31 | 0.78 | -1.21 – 1.83 | 0.40 | 0.691 | | | | | | |
| Positive | | | | | | | -0.38 | 0.60 | -1.56 – 0.80 | -0.63 | 0.529 | | | | | | |
| Autonomic | | | | | | | -0.62 | 0.84 | -2.26 – 1.02 | -0.74 | 0.459 | | | | | | |
| Individual | | | | | | | | | | | | | -0.77 | 0.50 | -1.75 – 0.21 | -1.54 | 0.123 |
| Interaction | | | | | | | | | | | | | -0.94 | 0.89 | -2.69 – 0.81 | -1.05 | 0.293 |
| Population | | | | | | | | | | | | | -2.84 | 0.79 | -4.39 – -1.29 | -3.58 | **<0.001** |
| Observations | 134 | | | | | | 147 | | | | | | 149 | | | | |
| $R^2$ Nagelkerke | 0.108 | | | | | | 0.011 | | | | | | 0.082 | | | | |

Table S2. Generalized linear model results testing for differences in residual-adjusted thermal optima among response categories. Separate Gamma GLMs were fit to (i) response type, (ii) response motivation, and (iii) level of biological organization. Intercepts represent the following reference categories: response type: metabolism; responsive motivation: negative;; response organization: internal ). Estimates, standard errors, and model fit statistics are reported.

| | *Response Type* | | | | | *Response Motivation* | | | | | *Response Organization* | | | |
|---|---|---|---|---|---|---|---|---|---|---|---|---|---|---|
| | **Activation Energy** | | | | | **Activation Energy** | | | | | **Activation Energy** | | | |
| *Predictors* | *Estimates* | *CI* | *Statistic* | *p* | | *Estimates* | *CI* | *Statistic* | *p* | | *Estimates* | *CI* | *Statistic* | *p* |
| Intercept (Metabolism) | 0.56 | 0.49 – 0.66 | -7.57 | **<0.001** | Intercept (Negative) | 0.47 | 0.38 – 0.59 | -6.68 | **<0.001** | Intercept (Internal) | 0.56 | 0.49 – 0.63 | -9.02 | **<0.001** |
| Energyaquisition | 1.11 | 0.80 – 1.60 | 0.61 | 0.541 | | | | | | | | | | |
| Somatic growth | 1.62 | 1.26 – 2.08 | 3.78 | **<0.001** | | | | | | | | | | |
| Locomotion | 0.88 | 0.65 – 1.21 | -0.82 | 0.414 | | | | | | | | | | |
| Reproduction | 2.24 | 1.54 – 3.39 | 4.03 | **<0.001** | | | | | | | | | | |
| Survival | 1.89 | 1.05 – 3.89 | 1.93 | 0.053 | | | | | | | | | | |
| Voluntary | | | | | | 1.10 | 0.75 – 1.65 | 0.48 | 0.632 | | | | | |
| Positive | | | | | | 1.36 | 0.92 – 2.08 | 1.50 | 0.134 | | | | | |
| Autonomic | | | | | | 1.71 | 1.31 – 2.21 | 4.04 | **<0.001** | | | | | |
| Individual | | | | | | | | | | | 1.38 | 1.13 – 1.70 | 3.07 | **0.002** |
| Interaction | | | | | | | | | | | 1.13 | 0.82 – 1.61 | 0.71 | 0.477 |
| Population | | | | | | | | | | | 2.19 | 1.58 – 3.11 | 4.54 | **<0.001** |
| Observations | 83 | | | | | 94 | | | | | 94 | | | |
| $R^2$ Nagelkerke | 0.359 | | | | | 0.225 | | | | | 0.255 | | | |

Table S3. Generalized linear model results testing for differences in activation energies among response categories. Separate Gamma GLMs with log link were fit for (i) response type, (ii) response motivation, and (iii) level of biological organization. Intercepts represent the following reference categories: response type: metabolism; response motivation: negative; response organization: internal. Estimates, standard errors, and model fit statistics are reported.

| | Activation Energy | | | | | Activation Energy | | | | |
|---|---|---|---|---|---|---|---|---|---|---|
| *Predictors* | *Estimates* | *std. Error* | *CI* | *Statistic* | *p* | *Estimates* | *std. Error* | *CI* | *Statistic* | *p* |
| (Intercept) | 1.44 | 0.21 | 1.02 – 1.87 | 6.94 | **<0.001** | 2.808 | 0.48 | 1.71 – 3.88 | 5.83 | **<0.001** |
| Performance Breadth | -0.07 | 0.02 | -0.12 – -0.03 | -23.20 | **0.003** | | | | | |
| Thermal Tolerance | | | | | | -0.08 | 0.02 | -0.13 – -0.03 | -3.50 | **0.007** |
| **Random Effects** | | | | | | | | | | |
| $\sigma^2$ | 0.02 | | | | | 0.05 | | | | |
| $\tau_{00}$ | 0.23 study_ID | | | | | 0.18 study_ID | | | | |
| ICC | 0.90 | | | | | 0.78 | | | | |
| N | 28 study_ID | | | | | 12 study_ID | | | | |
| Observations | 33 | | | | | 13 | | | | |
| Marginal $R^2$ / Conditional $R^2$ | 0.204 / 0.923 | | | | | 0.532 / 0.897 | | | | |

Table S4. Results of linear mixed-effects models testing relationships between activation energy and two measures of thermal breadth. Activation energy decreased significantly with both performance breadth and thermal tolerance breadth. Models included study ID as a random intercept to account for non-independence among observations from the same study. Estimates, standard errors, confidence intervals, test statistics, and p-values are shown.
